# Three-dimensional nanostructure of an intact microglia cell

**DOI:** 10.1101/388538

**Authors:** Giulia Bolasco, Laetitia Weinhard, Tom Boissonnet, Ralph Neujahr, Cornelius T. Gross

**Author notes:** Correspondence Dr. Cornelius Gross.

## Abstract

Microglia are non-neuronal cells of the myeloid lineage that invade and take up long-term residence in the brain during development (Ginhoux et al. 2010) and are increasingly implicated in neuronal maturation, homeostasis, and pathology (Bessis et al. 2007; Paolicelli et al. 2011; Li et al. 2012; Aguzzi et al. 2013, Cunningham 2013, Cunningham et al. 2013). Since the early twentieth century several methods for staining and visualizing microglia have been developed. Scientists in Ramón y Cajal’s group (Achúcarro 1913, Río-Hortega 1919) pioneered these methods and their work led to the christening of microglia as the *third element* of the nervous system, distinct from astrocytes and neurons. More recently, a combination of imaging, genetic, and immunological tools has been used to visualize microglia in living brain (Davalos et al. 2005; Nimmerjahn et al. 2005). It was found that microglia are highly motile under resting conditions and rapidly respond to injuries (Kettenmann et al. 2011) suggesting a role for microglia in both brain homeostasis and pathology. Transmission Electron microscopy (TEM) has provided crucial complementary information on microglia morphology and physiology but until recently EM analyses have been limited to single or limited serial section studies (Tremblay et al. 2010; Paolicelli et al. 2011; Schafer et al. 2012; Tremblay et al. 2012; Sipe et al. 2016). TEM studies were successful in defining a set of morphological criteria for microglia: a polygonal nucleus with peripheral condensed chromatin, a relatively small cytoplasm with abundant presence of rough endoplasmic reticulum (RER), and a large volume of lysosomes and inclusions in the perikaryon. Recent advances in volumetric electron microscopy techniques allow for 3D reconstruction of large samples at nanometer-resolution, thus opening up new avenues for the understanding of cell biology and architecture in intact tissues. At the same time, correlative light and electron microscopy (CLEM) techniques have been extended to 3D brain samples to help navigate and identify critical molecular landmarks within large EM volumes (Briggman and Denk 2006; Maco et al. 2013; Blazquez-Llorca et al. 2015, Bosch et al. 2015). Here we present the first volumetric ultrastructural reconstruction of an entire mouse hippocampal microglia using serial block face scanning electron microscopy (SBEM). Using CLEM we have ensured the inclusion of both large, small, and filopodial microglia processes. Segmentation of the dataset allowed us to carry out a comprehensive inventory of microglia cell structures, including vesicles, organelles, membrane protrusions, and processes. This study provides a reference that can serve as a data mining resource for investigating microglia cell biology.

## Material & Methods

### Confocal imaging & laser etching

*Thy1*::EGFP; *Cx3cr1*::CreER; *RC*::LSL-tdTomato triple transgenic mice were bred, genotyped and tested at EMBL following protocols approved by the EMBL Animal Use Committee and the Italian Ministry of Health. Mice were sacrificed at P15, perfused transcardially with PBS and fixed with Karnosky fixative (2% (w/v) PFA, 2.5% (w/v) Glutaraldehyde-TAAB) in 0.1 M Phosphate Buffer (PB). After perfusion brains were dissected, trimmed in *xy* around hippocampal areas and postfixed in 4% PFA in PB 0.1M overnight at 4°C. Subsequently, 60 µm thick vibratome (Leica Microsystems) coronal sections were cut and DAPI (Thermo Fisher) stained. Sections were mounted with 1% Low Melting Agarose (Sigma) in PB 0.1M on glass bottom dishes with alphanumeric grid (Ibidi). Regions of interest (ROI) containing microglia were imaged at low and high magnification with TCS SP5 resonant scanner confocal microscope (Leica Microsystems) with a 63x/1.2 water immersion objective, at a pixel size of 48 nm and a step size of 300 nm. Low magnification stacks containing the ROI were acquired in bright field, RFP, GFP and DAPI channels, thus creating as a navigation map with internal fiducial markers (microglia, capillaries and cell nuclei). A UV-diode laser operating at 405nm, a DPSS solid-state laser at 561 nm, and an Argon laser at 488 were used as excitation sources. Subsequent to confocal imaging, the external side of the glass-bottom grid was wet with 50% EtOH, carefully detached from the plastic dish and placed onto Laser Capture Micro-dissector (LMD7000, Leica Microsystems) for laser etching of the ROI.

### Serial Block Face Scanning Electron Microscopy (SBEM)

Laser branding sections were retrieved and stored in 0.5% PFA in PB 0.1M at 4°C. Selected sections were shaped in asymmetric polygons around the ROI and coordinates were measured before proceeding for electron microscopy (EM) preparation. Briefly sections were washed in cold Sodium Cacodylate buffer 0.1M pH 7.4, postfixed with 2% OsO_4_/1.5% Potassium Ferrocyanide for 1 h on ice, followed by a step with Thiocarbohydrazide for 20 min at RT and then a second step of 30 min in 2% aqueous OsO_4_ on ice. Samples were then rinsed carefully in water and stained “en block” first with 1% aqueous solution of Uranyl Acetate ON at 4°C and then with Lead Aspartate at 60°C for 30 min. Subsequently, sections were dehydrated with increasing concentration of Acetone and infiltrated in Durcupan resin overnight followed by 2 h embedding step with fresh resin. Durcupan embedding was carried out in a flat orientation within a sandwich of ACLAR^®^ 33C Films (Electron Microscopy Science) for 72h at 60°C. Upon embedding samples were trimmed using the coordinates to about 1 mm width to fit on the stab for 3View ultra-microtome (Gatan) and sputter coated with 5nm Iridium (EMITECH). Samples were mounted into aluminum pin using double component Epoxy resin supplemented with Carbon anno-tubes to provide electrical conductivity. In order to have a broader field of view (FOV), data were acquired on 3view fitted into a Gemini 330VP (Zeiss, Germany) SEM (scanning EM) column. Images were acquired first at low resolution (15*15 nm *xy* pixel size) and to reveal the etched marks from the laser micro-dissection, and correlate both vasculature and cell nuclei with the confocal imaging map. Upon pinpointing of the FOV containing the microglia of interest (31.72*23.80*22.6 μm) the imaging parameters were set to variable pressure with 5 Pa water vapor, 1.3kV HV high current, dwell time 6 µsec at a pixel size of 5 nm and section thickness 25 nm.

### Image processing and segmentation

A single stack file containing the 3view/GeminiSEM generated dataset was aligned using ImageJ software (Rasband and Bright 1995, https://imagej.nih.gov/ij/) with the plugin Linear Stack alignment with SIFT (Lowe 2004, https://imagej.net/Linear_Stack_Alignment_with_SIFT). To reduce dataset size and therefore further computation and memory usage, the stack was binned 2X in *xy* axis, to obtain a final *xy* resolution of 10×10 nm. At this resolution, it was still possible to identify the organelles and small processes. Microglia of interest was located based on its *xyz* coordinates within the ROI and from fiducial objects correlated between EM and confocal datasets. Segmentation of microglia cell body was carried out manually using iMOD software (Kremer et al. 1996, http://bio3d.colorado.edu/imod/) driven by a set of morphological criteria: the presence of abundant rough endoplasmic reticulum, the condensed chromatin at the periphery of the nucleus and the lack of mitochondria in the thin processes. A 3D model was then generated and correlated to the confocal dataset to confirm microglia identity. Systematic classification of organelles and vesicles trafficking within the cell was done by manual segmentation. Measurement of different color-code objects (Fig.2) was done by meshing the iMOD 3D model and exporting to the .obj format using built-in commands of iMOD (imodmesh.exe and imod2obj.exe). The generated mesh file was imported in Blender (https://www.blender.org/) to create and render in 3D an animation of the full EM dataset combined with the 3D model. Moreover, NeuroMorph, plugin (Jorstad et al. 2015, https://github.com/NeuroMorph-EPFL/NeuroMorph) was used to individually microglia’s main branches and filopodia.

**Figure 1.**
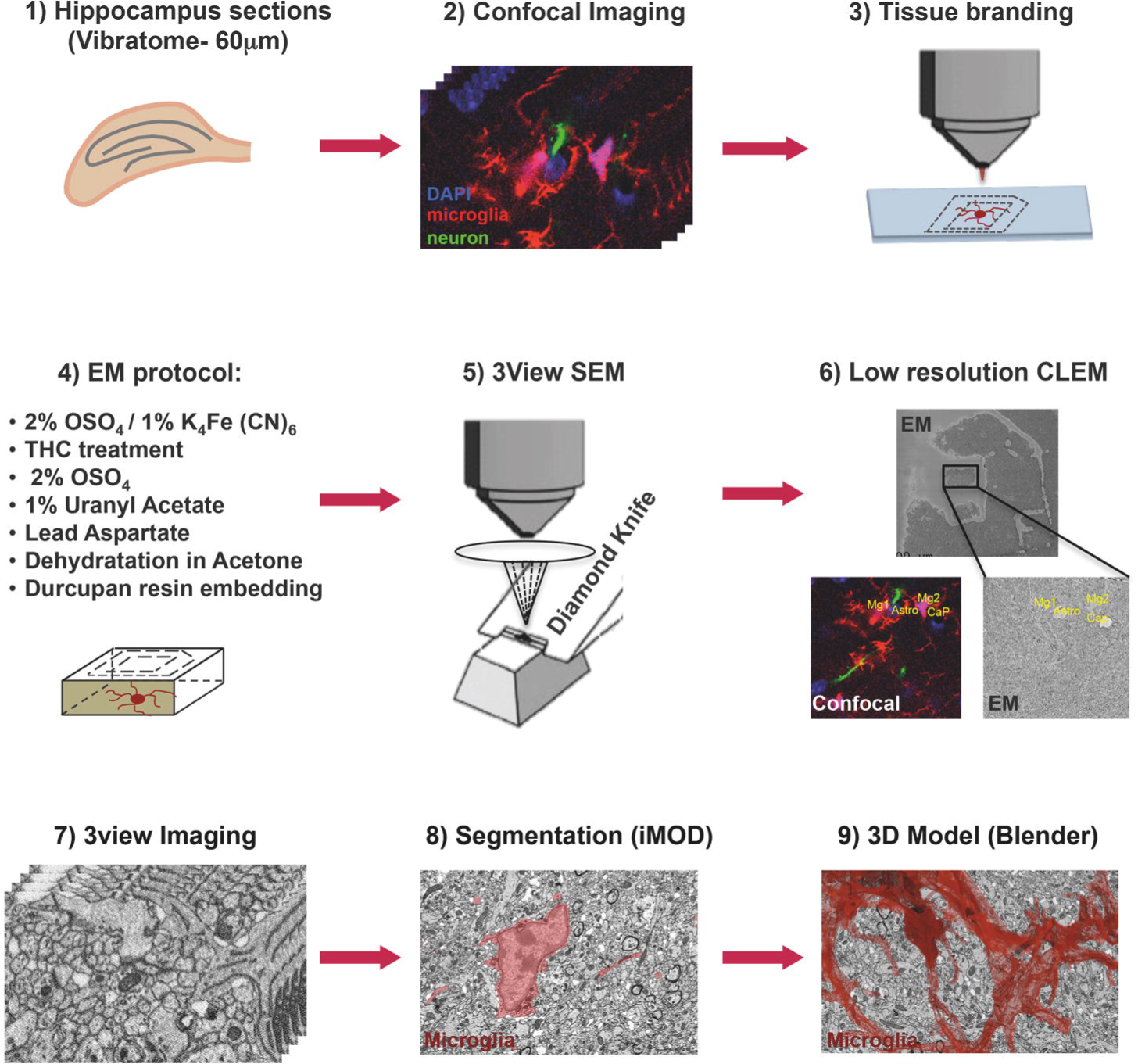
Correlative light and electron microscopy workflow. Confocal imaging of microglia (tdTomato, *red*), cell nuclei (DAPI, *blue*) and sparsely labeled neurons (EGFP, *green*), in sections from mouse hippocampus was followed by tissue laser branding, sample preparation for electron microscopy and serial block face EM, manual segmentation (iMOD), and rendering (Blender) of the microglia, *red*.

**Figure 2.**
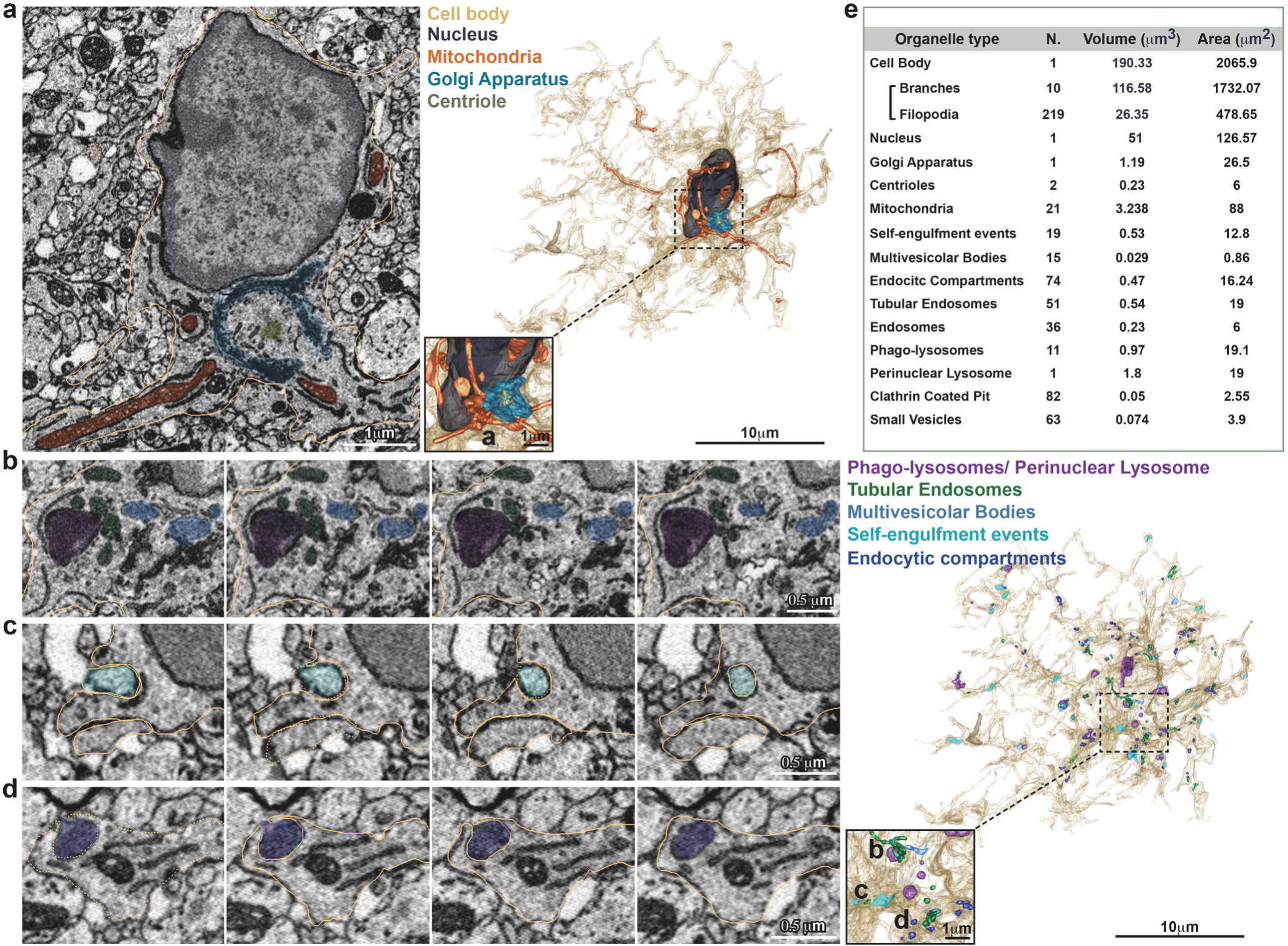
Inventory of microglia structures. Serial Block Face EM images showing segmentation of selected color-coded objects within the microglia (**a**-**d**, *gold*). **a**) Representative micrograph containing the microglia nucleus (*dark grey*), where also mitochondria (*orange*), Golgi apparatus (*indigo*) and centrioles (*grey*) are segmented (left). 3D surface rendering of peri-nuclear objects together with high magnification inset (right). **b-d**Representative series of consecutive micrographs showing the microglia: **b**) multi-vesicular bodies (*clear blue*) and phago-lysosomes (*purple*); **c**) self-engulfment of a branch originating from the same microglia (*turquoise*); **d**) early endocytic compartment (*dark blue*). 3D surface rendering of intra-cellular vesicles, together with high magnification inset (right). **e)**Total Volume and Area of segmented structures.

## Database description

The Serial Block Face EM series of stack has been split in 2 separate datasets due to the 4GB iMOD memory usage restrictions: -*dataset1*-(sect 1 to 389) and -*dataset2*-(section 390 to 904) respectively with their 3D models set at +/- 389 *z* offset. Each dataset collection is deposited at the Electron Microscopy Public Image Archive -EMPIAR- (Iudin et al. 2016) https://www.ebi.ac.uk/pdbe/emdb/empiar/entry/10201, and should be opened together with the related model using 3dmod on iMOD software, previously downloaded at http://bio3d.colorado.edu/imod/. Both a Zap and an Information window will be opened. The Information window has the main controls for black and white contrast, and allows for opening the Model View window by clicking on the *Image* button. The ZaP (Zoom and Pan) window contains the stack with the different color-coded segmented objects, and allows, by using the toolbar, several controls such as zooming in/out and sliding through the sections. In the Model View window objects can be switched on and off using the Edit button and the model can be rotated or edited. Additional features can be found at http://bio3d.colorado.edu/imod/doc/3dmodguide.html.

Moreover a -*Segmented.blend* file to visualize the stacks in *x*, *y*, *z* orientations integrated with its 3D Model can be found in the database (above link). The file can be opened with Blender (https://www.blender.org/) and can be explored through upon installation of the provided add-on -NeuroMorph_3D_Drawing.py- (see NeuroMorph documentation for help). Additional features are available with the complete NeuroMorph add-ons suite (https://github.com/NeuroMorph-EPFL/NeuroMorph).

To quickly present the dataset and show the rendering capabilities of Blender, a 3D animation -*Blendervideo.avi*- has also been stored in the repository. It shows the full SBEM dataset in the 3 orthogonal anisotropic planes meshed with the 3D model of few selected objects (microglia cell body-*dark grey*, nucleus-*blue*, mitochondria-*orange* and lyso-phagosomes-*purple*).

## Author Contributions

C.T.G. conceived the project. G.B. and L.W. set up the protocol and prepared the samples. G.B. segmented the full dataset and wrote the paper. R.N. ran the 3View SBEM system. T.B. processed and measured the 3D model and created the animation.

## Acknowledgements

Funding has been provided by EMBL (C.T.G., G.B., L.W., and T.B.) and ERC Advanced Grant COREFEAR (C.T.G),

